# Applying computational protein design to engineer affibodies for affinity-controlled delivery of vascular endothelial growth factor and platelet-derived growth factor

**DOI:** 10.1101/2024.12.18.629300

**Authors:** Justin E. Svendsen, Madeleine R. Ford, Chandler L. Asnes, Simon C. Oh, Jonathan Dorogin, Karly M. Fear, Johnathan R. O’Hara-Smith, Michael J. Harms, Parisa Hosseinzadeh, Marian H. Hettiaratchi

**Affiliations:** Knight Campus for Accelerating Scientific Impact, University of Oregon; Department of Chemistry and Biochemistry, University of Oregon; Institute of Molecular Biology, University of Oregon; Department of Human Physiology, University of Oregon; Department of Biology, University of Oregon

## Abstract

Effective angiogenesis requires the coordinated presentation of vascular endothelial growth factor (VEGF) and platelet-derived growth factor (PDGF), which play competing roles in stimulating vascular outgrowth and stabilization. Dysregulation in VEGF and PDGF secretion are implicated in abnormal vascular structures, poor vessel stability, and inadequate tissue repair. Current biomaterial delivery vehicles for these proteins have a limited ability to precisely control the kinetics of protein release, preventing systematic exploration of their temporal effects. Here, we combined yeast surface display and computational protein design to engineer eight VEGF-specific and PDGF-specific protein binders called affibodies with a broad range of affinities (dissociation constants = 2.23-9260 nM) for affinity-controlled protein release. Starting from one VEGF- and one PDGF-specific affibody discovered from yeast surface display, computational modeling was used to select disruptive mutations to create VEGF-specific affibodies with lower affinities for VEGF and expand the affinity range of PDGF-specific affibodies. In both cases, the specificity of the affibodies for their target protein was either maintained or enhanced. Soluble protein-specific affibodies modulated protein bioactivity as evidenced by changes in VEGF-induced endothelial cell proliferation and luminescent output of a PDGF-responsive fibroblast cell line. Affibody-conjugated polyethylene glycol maleimide (PEG-mal) hydrogels sustained VEGF and PDGF release compared to hydrogels without affibodies, enabling tunability of protein release over 7 days. VEGF and PDGF released from affibody-conjugated hydrogels exhibited higher bioactivity compared to proteins released from hydrogels without affibodies, highlighting the potential of these engineered affinity interactions to prolong protein bioactivity. This work underscores the power of computational protein design to enhance biomaterial functionality, creating a platform for tunable protein delivery to support angiogenesis and tissue repair.

## Introduction

Coordinated secretion of multiple proteins is required for both tissue development and repair.^1,2^ In the case of angiogenesis after injury, the expansion of existing vascular networks requires a variety of morphogens, including vascular endothelial growth factors (VEGF) and platelet derived growth factors (PDGF), which are secreted by fibroblasts, macrophages, endothelial cells, and other support cells proximal to the injury site.^3–5^ VEGF destabilizes pericyte-endothelial cell contacts, transforms endothelial cells into motile tip cells, and stimulates tip cell migration toward the injury site.^6,7^ Newly differentiated stalk cells secrete PDGF that stimulates pericyte adherence to the endothelial cell wall for vessel stabilization, resulting in the downstream formation of mature vasculature.^5,8,9^ The contrasting roles of VEGF and PDGF in stimulating and stabilizing vascular outgrowth require their carefully regulated balance *in vivo*.^10–15^ Dysregulation in VEGF and PDGF secretion within injured tissues can cause aberrant vascular geometries, poor vessel stability, and inadequate tissue coverage, highlighting the need for therapeutic interventions that can restore normal VEGF and PDGF concentrations after injury.^12,16^ As such, sustained delivery of exogenous VEGF and PDGF to injured tissues holds potential for re-establishing vascular homeostasis.^17^

Several biomaterial-based protein delivery vehicles have enabled sustained delivery of VEGF and PDGF and improved angiogenesis through diffusion-controlled protein release.^18–20^ However, current delivery vehicles have a limited ability to control the kinetics of protein release in injury environments, in which protein concentrations rapidly change, preventing systematic exploration of the effects of temporal protein delivery on angiogenic cell signaling.^21^ To address these limitations, biomaterials that harness specific, reversible protein-material affinity interactions have emerged as a promising method to finely tune protein presentation and bioactivity.^1,22^ Extracellular matrix (ECM) molecules, such as collagen, fibronectin, and heparin, that naturally engage in affinity interactions with heparin-binding proteins in the body have been incorporated into biomaterials delivery vehicles to control protein release.^23–26^ However, since the angiogenic isoforms of VEGF (VEGF165) and PDGF (PDGF-BB) both contain ECM-binding domains and share structural similarities,^27–29^ independent control over the release of these proteins requires the development of highly specific protein-material interactions.

Current approaches for developing specific protein-protein interactions for affinity-controlled protein release leverage directed evolution and cell sorting of phage and yeast display libraries of protein binders.^30–35^ These strategies can be time-intensive and are restricted by library sequence diversity and scaffold structure, limiting the success of binder discovery and control over the characteristics of resulting protein-protein interactions.^36,37^ Alternatively, recent developments in computational approaches for rational protein design have enabled the design of protein-protein interactions with user-defined characteristics beyond the reach of directed evolution, accelerating the discovery of protein binders.^38,39^ Thus, we sought to combine the advantages of yeast surface display and rational protein design to engineer highly specific protein-protein affinity interactions for VEGF165 and PDGF-BB across a broad range of affinities with the goal of tuning protein release and bioactivity from biomaterials to stimulate angiogenesis.

Here, we report a new collection of variable affinity VEGF- and PDGF-specific affibodies, small alpha-helical protein binders, that enable temporal control over protein release and bioactivity. We demonstrate that VEGF- and PDGF-specific affibodies identified using yeast surface display can be diversified using rational protein design without losing protein specificity. We used computational modeling to inform the selection of disruptive point mutations to a high-affinity VEGF-specific affibody, resulting in three mutants with lower affinities for VEGF. To expand the affinity range of our low-affinity PDGF-specific affibody, we used Rosetta-based rational design to engineer three PDGF-specific affibody mutants with different affinities for PDGF. We demonstrated that VEGF- and PDGF-specific affibodies conjugated to polyethylene glycol maleimide (PEG-mal) hydrogels controlled VEGF and PDGF release based on their binding affinities. Soluble VEGF-specific and PDGF-specific affibodies modulated the bioactivity of their respective proteins as determined by VEGF-induced proliferation of human umbilical vein endothelial cells (HUVECs) and luminescent output of a PDGF-responsive fibroblast cell line. VEGF and PDGF released from affibody-conjugated hydrogels displayed higher bioactivity than protein released from PEG-mal hydrogels without affibodies, suggesting that affinity interactions between proteins and affibodies may prolong protein bioactivity. Overall, this work establishes a library of novel VEGF- and PDGF-specific binders capable of precisely controlling the release of bioactive VEGF and PDGF from hydrogels, while demonstrating the successful use of computational protein design to engineer protein binders with high specificity and diverse affinities from known starting scaffolds.

## Results and Discussion

### Yeast Surface Display Identifies Novel Protein Binders for VEGF and PDGF

We sought to identify affibodies that could specifically bind to either VEGF165 or PDGF-BB by applying iterative rounds of magnetic-activated cell sorting (MACS) and fluorescent-activated cell sorting (FACS) to a yeast surface display library of approximately 400 million affibody variants.^30,34^ We performed four rounds of MACS using magnetic beads covalently conjugated with either VEGF165 or PDGF-BB as the positive sort target, while bovine serum albumin (BSA)-conjugated beads and Tris-coated beads were used for negative sorting.^34^ Following MACS, we performed two rounds of FACS to enrich the population for affibody-expressing yeast that bound to the target protein. We selected one VEGF-specific affibody and one PDGF-specific affibody identified using this method for further analysis.

### Computational Modeling Predicts Point Mutations that Alter VEGF Affibody Binding Affinity

Yeast displaying the VEGF-specific affibody were incubated with a range of biotinylated VEGF concentrations (bVEGF165; 0.02-10 µM) to determine an equilibrium dissociation constant (K_D_) of 861 ± 255 nM using flow cytometry (**Fig. 1A**). This affinity between VEGF and the VEGF-specific affibody is similar to that of known affinity interactions between VEGF165 and heparin (K_D_ = 165 nM) and that of heparin with several other proteins involved in angiogenesis, such as integrin α5β1 (K_D_ = 2040 nM), antithrombin-III (K_D_ = 163 nM), and fibroblast growth factor-1 (K_D_ = 461 nM).^40^

**Figure 1.**
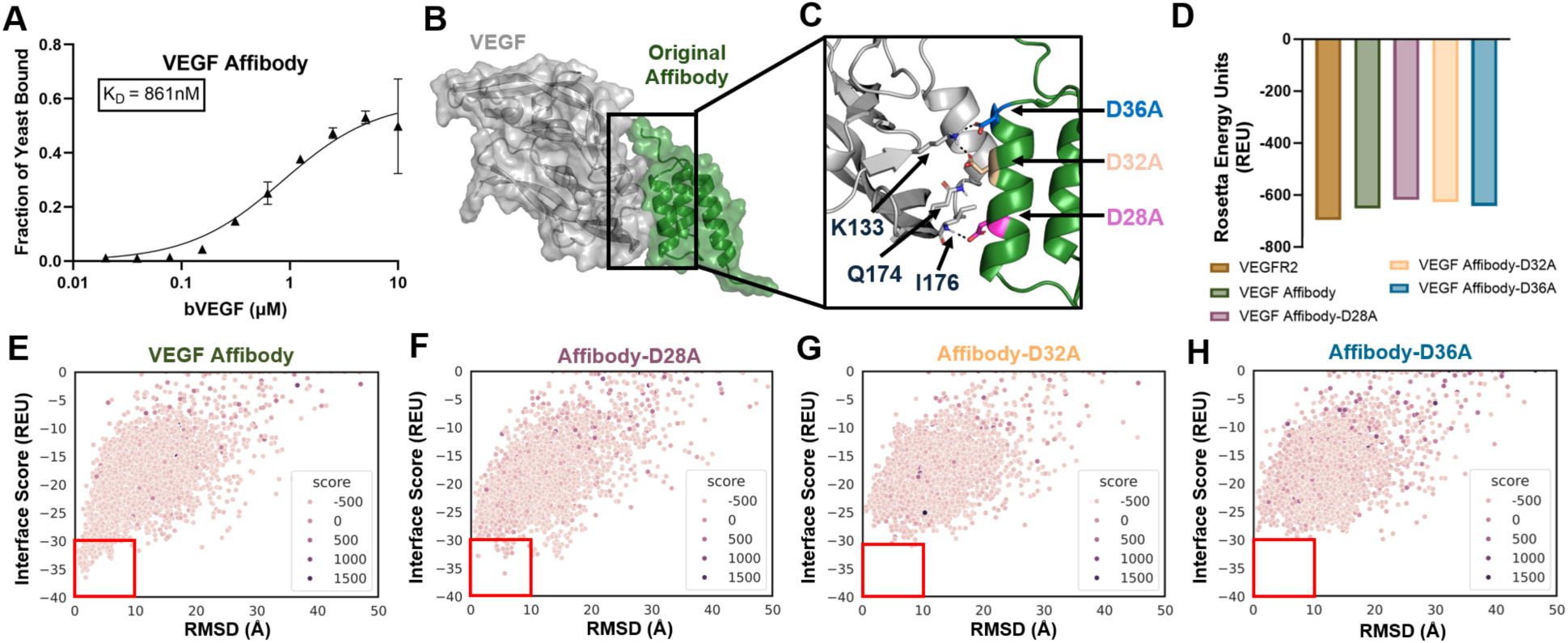
Computational modeling of VEGF-specific affibodies identifies destabilizing mutations and generates novel VEGF affibody mutants. A) Flow cytometry analysis of fraction of VEGF-specific affibody-displaying yeast binding to 0.02-10 µM of bVEGF, exhibiting an affinity interaction of K_D_ = 861 ± 255 nM (n=3). B) Computational molecular modeling of the binding interface between VEGF (grey) and the original VEGF affibody (green). C) Inset depicting key residues on VEGF and the VEGF-specific affibody interacting at the VEGF-affibody binding interface. VEGF affibody residues selected for mutagenesis include D28A (pink), D32A (beige), and D36A (blue). D) Rosetta Scores, shown as Rosetta Energy Units, comparing interface stabilities for VEGFR-2, the original VEGF affibody, VEGF Affibody-D28A, VEGF Affibody-D32A, and VEGF Affibody-D36A binding to VEGF at its VEGFR-2 binding epitope. Rosetta docking funnels measuring the interface stabilities of VEGF-affibody binding across the entire surface of VEGF for E) the original VEGF affibody, F) VEGF Affibody-D28A, G) VEGF Affibody D32A, and H) VEGF Affibody D36A, with red boxes highlighting binding interactions close to the intended site.

Given that VEGF is critical in the early stages of angiogenesis, we wanted to engineer additional affibodies that would allow us to create biomaterials that rapidly released VEGF. We set out to identify additional protein binders with lower affinities for VEGF. We used ZDOCK, HDOCK, AlphaFold2, and Rosetta protein design softwares to identify point mutations with a high likelihood of decreasing the affinity strength of the original VEGF-affibody interaction while maintaining the specificity of the affibody for VEGF. We generated a structure for the VEGF-specific affibody from yeast surface display using Alphafold2 and a structure of VEGF from its x-ray crystallography data (PDB: 2VPF). To establish a computational reference for a known binding interaction with VEGF, we generated a structure of VEGF bound to VEGF receptor-2 (VEGFR-2) by combining elements from the VEGF crystal structure 2VPF with the VEGF-VEGFR-2 crystal structure 3V2A.^41^ Rosetta FastRelax was used to generate the lowest energy conformation of surface-exposed residues,^42^ and computational docking with ZDOCK and HDOCK^43,44^ was used to predict the potential binding interfaces between VEGF and the VEGF-specific affibody. Rosetta Score, which is negatively correlated with thermodynamic stability, was used to evaluate the stability of these binding interfaces.^45,46^

These analyses indicated the most probable VEGF-affibody binding interface to be at the VEGFR-2 binding epitope on VEGF (**Fig. 1B, Fig. S1A**). Within this interface, three aspartic acids on the VEGF affibody at positions 28, 32, and 36 were predicted to participate in polar contacts with residues K133, M166, and I176 on VEGF, contributing to the formation of a hydrophobic pocket at the core of the protein-protein interaction **(Fig. 1C)**. We individually mutated these three aspartic acids on the affibody to alanine and re-calculated the docking scores to determine the predicted impact on the binding stability, creating three point mutants we named VEGF Affibody-D28A, VEGF Affibody-D32A, and VEGF Affibody-D36A.^47,48^

### Computational Analysis of VEGF-specific Affibodies Predicts Binding to the VEGFR-2 Epitope on VEGF

Rosetta Scores for predicted interactions between VEGF and the mutant affibodies at the VEGFR-2 binding epitope were higher and less stable than predicted interactions between VEGF and the original VEGF affibody, supporting the hypothesis that each point mutation would disrupt VEGF-affibody binding (**Fig. 1D**). Rosetta Docking algorithms using the VEGFR-2 epitope on VEGF as the starting pose indicated a single stable binding interface at this location for all affibodies (**Fig. 1E-H**). Higher interface scores were also observed during docking for all three VEGF affibody mutants compared to the original VEGF affibody. As expected, the known VEGF-VEGFR-2 binding interaction demonstrated the lowest Rosetta Score and interface scores in Rosetta Docking (**Fig. 1D, S1B**), indicating the highest stability and suggesting the VEGF affibodies would bind to VEGF with weaker affinity than VEGFR-2.

### Computational Modeling Suggests VEGF-specific Affibodies Will Not Interact with PDGF

Rosetta Scores for VEGF-specific affibodies binding to PDGF at the PDGF receptor-beta (PDGFRβ) epitope were higher than that of VEGF-specific affibodies binding to VEGF (**Fig. S1C**). Rosetta Docking predicted no single stable interface for any VEGF-specific affibody binding to PDGF (**Fig. S1D**). These *in silico* results suggested that the VEGF affibodies would be unlikely to bind to PDGF.

### Rosetta-based Rational Design of PDGF-specific Affibodies

Yeast displaying the PDGF-specific affibody were incubated with a range of biotinylated PDGF concentrations (bPDGF-BB; 0.016-8.4 µM), and PDGF binding to yeast was analyzed via flow cytometry to determine an equilibrium dissociation constant of 855 ± 238 nM (**Fig. 2A**). Similar to VEGF, we sought to engineer additional affibodies with a broader range of binding affinities for PDGF to tune protein release. However, because PDGF typically plays a role in the later stages of angiogenesis,^5,49^ we aimed to engineer PDGF binding affibodies with higher binding affinities, resulting in slower, sustained protein release. To achieve this, we used Rosetta to rationally mutate residues on the PDGF-specific affibody that would increase its affinity for PDGF. To model the PDGF-affibody binding interaction, we generated a structure for the PDGF-specific affibody using Alphafold2 and derived a structure of PDGF from the x-ray crystallography data of PDGF bound to PDGFRβ (PDB: 3MJG).^50^ This crystal structure of PDGF bound to PDGFRβ was also used as a known binding interaction to which computational results could be compared.

**Figure 2.**
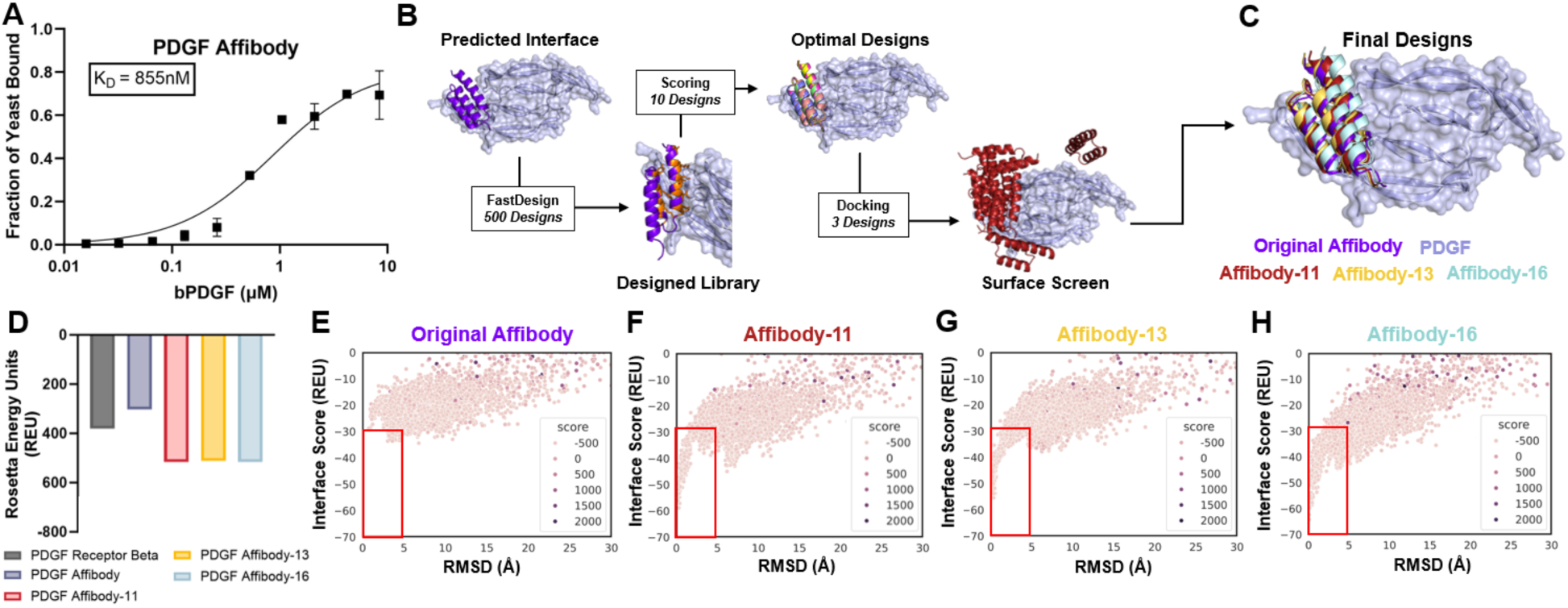
Rosetta-based computational design generates novel PDGF-specific affibodies with high-stability binding interactions with PDGF. A) Flow cytometry analysis of fraction of PDGF-specific affibody-displaying yeast binding to 0.016-8.4 µM of bPDGF, exhibiting an affinity interaction of K_D_ = 855 ± 238 nM (n = 3). B) Rosetta computational design pipeline for engineering mutant PDGF-specific affibodies from the original PDGF-specific affibody. C) Computational modeling of the binding interfaces between PDGF (light purple) and the original PDGF-specific affibody (dark purple), PDGF Affibody-11 (red), PDGF Affibody-13 (yellow), and PDGF Affibody-16 (light blue). D) Rosetta Scores comparing interface stabilities for PDGFRβ, the original PDGF Affibody, PDGF Affibody-11, PDGF Affibody-13, and PDGF Affibody-16 binding to PDGF at its PDGFRβ binding epitope. Rosetta Docking funnels measuring the interface stabilities of PDGF-affibody binding across the entire surface of PDGF for E) the original PDGF Affibody, F) PDGF Affibody-11, G) PDGF Affibody-13, H) PDGF Affibody-16, with red boxes highlighting binding interactions close to the intended site.

Rosetta FastRelax was used to generate the lowest energy conformation of the surface-exposed residues on PDGF and the PDGF-specific affibody,^42^ and ZDOCK and HDOCK were used to generate potential binding interfaces of the PDGF-affibody binding interaction. Rosetta Scoring and comparisons to known receptor binding interfaces predicted the most stable binding interface to overlap with the PDGFRβ binding epitope (**Fig. 2B**, **Fig. S2A**). We then generated 500 mutant affibodies using Rosetta FastDesign,^51^ by allowing mutagenesis of affibody residues in direct contact with the predicted PDGF binding interface to any amino acid except cysteine to avoid the introduction of disulfide bridges. Structures of mutant affibodies bound to PDGF were screened using Rosetta-generated score metrics to select for interfaces with high shape complementarity, low solvent accessible surface area, high hydrophobic contact surface area, no buried unsatisfied polar contacts, and high total interface contact surface area. This analysis resulted in 10 affibodies that were predicted to form stable binding interactions with PDGF.^39,45^ Further screening with Rosetta Docking algorithms using the PDGFRβ epitope on PDGF as the starting pose yielded three affibodies that were predicted to bind at the PDGFRβ epitope with high interface stability **(Fig. 2C)**. Next, we used Alphafold2 and Robetta folding packages to confirm that the *in silico* mutagenesis was unlikely to disrupt the alpha-helical folding of the designed affibodies.^52,53^ We named these affibodies PDGF Affibody-11, PDGF Affibody-13, and PDGF Affibody-16 to indicate the number of mutations from the original PDGF-specific affibody.

The Rosetta Scores of the three mutant PDGF-specific affibodies binding to PDGF at the PDGFRβ binding epitope were lower than the Rosetta Scores of the original PDGF affibody and PDGFRβ binding to PDGF, suggesting an increased likelihood of the mutant affibodies to stably bind to the PDGFRβ epitope on PDGF (**Fig. 2D**). While Rosetta Docking analysis of the original PDGF affibody binding to PDGF did not indicate a single stable binding interface within 10 Å of the PDGFRβ binding epitope (**Fig. 2E**), suggesting a low likelihood of successful binding to this designated site, Rosetta Docking algorithms predicted low interface scores at low root mean squared deviation (RMSD) from the PDGFRβ binding epitope for all mutant PDGF-specific affibodies (**Fig. 2F-H**) that were comparable to interface scores of the known PDGF-PDGFRβ binding interaction (**Fig. S2B**).

### Computational Modeling Suggests PDGF-specific Affibodies Will Not Interact with VEGF

Rosetta Scores of PDGF-specific affibodies binding to VEGF at the VEGFR-2 epitope were higher than that of PDGF-specific affibodies binding to PDGF (**Fig. S2C**). Rosetta Docking algorithms similarly revealed no single stable interface for binding between VEGF and the PDGF-specific affibodies (**Fig. S2D**), indicating a low likelihood of PDGF-specific affibodies non-specifically binding to VEGF.

### Biochemical Characterization of VEGF-specific and PDGF-specific Affibodies

VEGF and PDGF affibodies were recombinantly expressed in BL21 (DE3) *Escherichia coli* and purified from sonicated bacterial lysate using immobilized metal affinity chromatography followed by size exclusion chromatography. We obtained high-purity soluble affibodies at the expected molecular weight of 7 kDa, which was confirmed by sodium dodecyl sulfate-polyacrylamide gel electrophoresis (SDS-PAGE) (**Fig. S3**) and mass spectrometry (**Fig. S4A, S4B**). All affibodies displayed far-UV circular dichroism spectra consistent with their predicted alpha-helical structure (**Fig. S4C-D**).^54^

### Computationally Informed Single Point Mutations Decreased VEGF-specific Affibody Affinity for VEGF

Biolayer interferometry (BLI) was used to determine the binding kinetics and binding specificity of VEGF-specific and PDGF-specific affibodies for their respective proteins. Streptavidin-coated BLI probes were loaded with 25 nM of bVEGF, followed by association of 31.25-1000 nM of each of the VEGF affibodies (**Fig. S5A**). The equilibrium dissociation constant measured by BLI for the original VEGF affibody binding to VEGF (K_D_ = 89.8 nM) (**Table. 1**, **Fig. S5B**) was an order of magnitude lower than the dissociation constant measured via flow cytometry of the affibody-displaying yeast (K_D_ = 861 ± 255 nM) (**Fig. 1A**). The weaker affinity observed on yeast surface display was likely due to increased steric hindrance and decreased mobility of the VEGF-affibody binding interface when the affibody was fixed within the yeast surface display construct compared to being freely accessible in solution during BLI.

**Table 1.** Kinetic constants of VEGF-specific and PDGF-specific affibodies binding to VEGF and PDGF measured by biolayer interferometry. Binding between VEGF and VEGF-specific affibodies or PDGF and PDGF-specific affibodies was measured using biolayer interferometry. Data were globally fit to calculate equilibrium dissociation constants (K_D_), association rate constants (k_off_), and dissociation rate constants (k_on_). 95% confidence intervals are reported for each kinetic constant.

| Affibody | $k_{\text{off}} (\text{s}^{-1})$ | $k_{\text{off}} (\text{s}^{-1})$ 95% CI | $k_{\text{on}} (\text{M}^{-1} \text{s}^{-1})$ | $k_{\text{on}} (\text{M}^{-1} \text{s}^{-1})$ 95% CI | $K_D (\text{nM})$ |
| --- | --- | --- | --- | --- | --- |
| VEGF Affibody | 0.003861 | (0.003752, 0.003980) | 42,970 | (41,764, 44,162) | 89.8 |
| VEGF Affibody-D28A | 0.004180 | (0.004144, 0.004219) | 9190 | (8911, 9468) | 455 |
| VEGF Affibody-D32A | 0.00769 | (0.00709, 0.00770) | 10.8 | (9.1, 12.4) | 984,000 |
| VEGF Affibody-D36A | 0.008368 | (0.008308, 0.008421) | 4080 | (3900, 4260) | 2050 |
| PDGF Affibody | 0.000592 | (0.000589, 0.000594) | 180,220 | (179,559, 180,861) | 3.28 |
| PDGF Affibody-11 | 0.003107 | (0.003044, 0.003174) | 476,180 | (475,493, 489,419) | 6.44 |
| PDGF Affibody-13 | 0.01822 | (0.01756, 0.01892) | 236,620 | (225,424, 246,158) | 77.4 |
| PDGF Affibody-16 | 0.002643 | (0.002590, 0.002686) | 449,710 | (444,311, 455,173) | 5.87 |

We next turned our attention to the VEGF affibody mutants. All VEGF affibody mutants displayed weakened affinity towards VEGF (K_D_^D28A^ = 455 nM; K_D_^D32A^ = 984,000 nM; K_D_^D36A^ = 2050 nM) (**Table. 1; Fig. S5C-E**) compared to the original VEGF-specific affibody. This was consistent with the computationally predicted effect of introducing unsatisfied polar contacts at the VEGF-affibody binding interface. Comparatively, the VEGF Affibody-D36A and VEGF Affibody-D28A displayed moderate reductions in affinity for VEGF, likely due to neighboring residue side chains permitting repacking of the binding interface to interact with other solvent-exposed polar residues or water molecules. VEGF Affibody-D32A and VEGF Affibody-D36A displayed higher dissociation rate constants (k_off_) than the original VEGF affibody and VEGF Affibody-D28A, with all dissociation rate constants measured within the same order of magnitude (**Table. 1**). In contrast, all mutant VEGF-specific affibodies displayed lower association rate constants than the original VEGF affibody, spanning three orders of magnitude (**Table. 1**). The dramatic changes in association rate constants coupled with smaller changes in dissociation rate constants suggest that the point mutations mainly impacted recognition and binding to VEGF rather than its release.^55^ None of the VEGF-specific affibodies displayed off-target binding to PDGF despite its structural similarities to VEGF (**Fig. S5F**), confirming our Rosetta predictions and demonstrating that our computationally informed single point mutations could alter the affinity of the VEGF-specific affibody without compromising its specificity.

### Rosetta Rational Design Generates PDGF-specific Affibodies with Unique Binding Kinetics

Binding of PDGF to PDGF-specific affibodies was evaluated by loading nickel nitrilotriacetic acid (Ni-NTA)-coated BLI probes with 200 nM of each of the PDGF affibodies, followed by association of 1.563-50 nM of PDGF (**Fig. S5G**). The equilibrium dissociation constant of the original PDGF-specific affibody binding to PDGF was two orders of magnitude lower on BLI (K_D_ = 3.28 nM) than the equilibrium dissociation constant measured by flow cytometry (K_D_ = 855 ± 238 nM) **(Table. 1, Fig. S5H)**. Similar to the VEGF-specific affibody, the lower observed affinity between the original PDGF-specific affibody and PDGF on yeast surface display could be due to increased steric hindrance and decreased mobility of the PDGF-affibody binding interface compared to BLI.

PDGF Affibody-11 and PDGF Affibody-16 displayed similar affinities for PDGF compared to the original PDGF affibody (K_D_^Affibody-11^ = 6.44 nM, K_D_^Affibody-16^ = 5.86 nM), while PDGF Affibody-13 displayed an affinity that was an order of magnitude lower (K_D_^Affibody-13^ = 77.35 nM) (**Table. 1, Fig. S5I-K**). Except for the dissociation rate constant of PDGF Affibody-13, the association and dissociation rate constants of all the PDGF-specific affibodies measured within the same order of magnitude (**Table. 1**), with PDGF Affibody-11 and Affibody-16 displaying faster association rate constants than the original PDGF affibody. PDGF Affibody-13 displayed both a higher equilibrium dissociation constant and higher dissociation rate constant, demonstrating that rational design changed the kinetics of affibody binding instead of only increasing the affinity as intended. While the original PDGF-specific affibody displayed some off-target binding to VEGF, none of the mutant PDGF affibodies displayed binding to VEGF (**Fig. S5L**), suggesting that rational design was successful in increasing the specificity of the PDGF-affibody interaction.

### VEGF-specific and PDGF-specific Affibodies Control VEGF and PDGF Release from PEG Hydrogels

We next tested whether our designed affibodies could tune the kinetics of VEGF and PDGF release from hydrogels. Affibody-conjugated polyethylene glycol maleimide (PEG-mal) hydrogels were synthesized as previously described to contain 500 molar excess of either VEGF-specific or PDGF-specific affibodies and loaded with 100 ng of their respective protein overnight.^34^ Following removal of unencapsulated VEGF and PDGF, hydrogels were incubated in 0.1% of bovine serum albumin (BSA) in phosphate-buffered saline (PBS) and allowed to release protein over 7 days at 37 °C.

Hydrogels conjugated with VEGF Affibody-D28A encapsulated more VEGF than hydrogels without affibodies, while all other hydrogels displayed comparable protein encapsulation (**Fig. 3A**). Furthermore, hydrogels containing VEGF Affibody-D28A released less VEGF over 7 days than hydrogels containing VEGF Affibody-D32A or hydrogels without affibodies (**Fig. 3B**), with differences observed as early as 15 minutes (**Fig. 3C**). At intermediate time points, hydrogels conjugated with VEGF Affibody-D36A demonstrated lower VEGF release than hydrogels without affibodies and hydrogels containing VEGF Affibody-D32A, while displaying higher VEGF release than hydrogels conjugated with VEGF Affibody-D28A and the original VEGF affibody. These release profiles were consistent with the BLI data that demonstrated a 10,000-fold reduction in VEGF Affibody-D32A’s affinity for VEGF and a 20-fold reduction in VEGF Affibody-D36A’s affinity for VEGF compared to the original VEGF affibody. Thus, mutagenesis weakened the original VEGF-affibody affinity interaction without completely disrupting its ability to control VEGF release.

**Figure 3.**
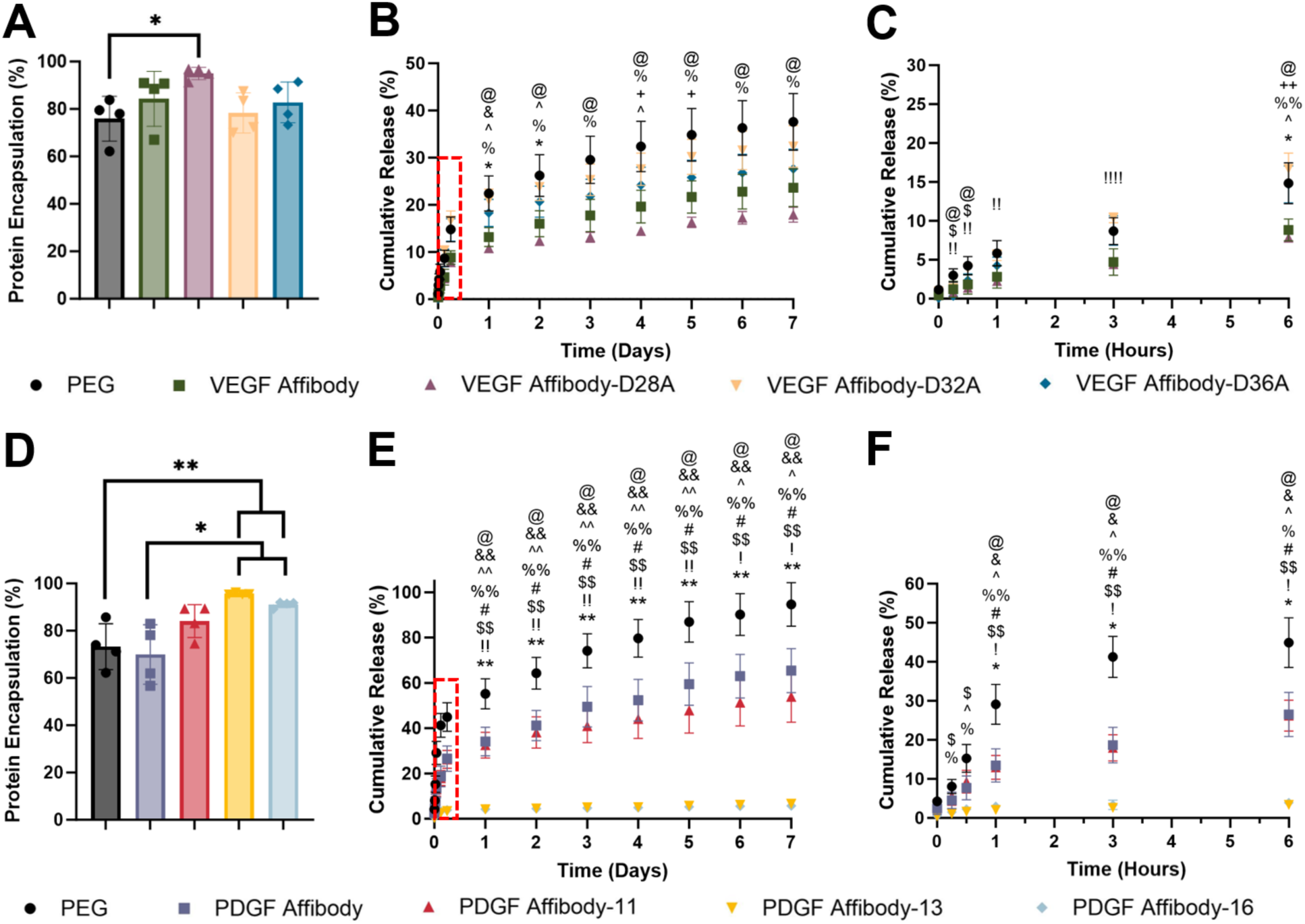
Protein release from VEGF-specific and PDGF-specific affibody-conjugated hydrogels. A) Encapsulation of VEGF within VEGF affibody-conjugated PEG-mal hydrogels. Statistical significance was determined by one-way ANOVA and Tukey’s post-hoc test. (n = 4, *p < 0.05) B) Cumulative release of VEGF from VEGF affibody-conjugated hydrogels at 37°C over 7 days measured by ELISA. C) Zoomed in graph of the first 6 hours of VEGF release indicated by dotted red box. Statistical significance was determined by two-way ANOVA and Tukey’s post-hoc test. (n = 4, one symbol = p < 0.05, two symbols = p < 0.01, and four symbols = p < 0.0001 as indicated. @ PEG vs. D28A, & PEG vs. VEGF Affibody, ^ D28A vs. D36A, % D28A vs. D32A, * VEGF Affibody vs. D32A, # PEG vs. D32A, $ PEG vs. D36A, ! D32A vs. D36A, + VEGF Affibody vs. D36A) D) Encapsulation of PDGF within PDGF affibody-conjugated PEG-mal hydrogels. Statistical significance was determined by one-way ANOVA and Tukey’s post-hoc test. (n = 4, *p < 0.05, **p < 0.01) E) Cumulative release of PDGF from PDGF affibody-conjugated hydrogels at 37°C over 7 days measured by ELISA. F) Zoomed in graph of the first 6 hours of PDGF release indicated by dotted red box. Statistical significance was determined by two-way ANOVA and Tukey’s post-hoc test. (n = 4, one symbol = p < 0.05 and two symbols = p < 0.01 as indicated. @ PEG vs. PDGF Affibody, & PDGF Affibody vs. Affibody-13, ^ Affibody-11 vs. Affibody-13, % PEG vs. Affibody-16, # PEG vs. Affibody-11, $ PEG vs. Affibody-13, ! Affibody-11 vs. Affibody-16, * PDGF Affibody vs. Affibody-16)

Hydrogels conjugated with either PDGF Affibody-13 or PDGF Affibody-16 encapsulated more PDGF than hydrogels conjugated with the original PDGF affibody and hydrogels without affibodies **(Fig. 3D)**. All hydrogels containing affibodies displayed reduced PDGF release after 7 days compared to hydrogels without affibodies,with identical differences between groups for Days 1-7 **(Fig. 3E).** Hydrogels containing PDGF Affibody-13 and PDGF Affibody-16 further reduced PDGF release over 7 days compared to all other groups with differences in release profiles observed as early as 1 and 3 hours, respectively **(Fig. 3F)**. Despite the similar PDGF release profiles observed for hydrogels containing PDGF Affibody-13 and PDGF Affibody-16, PDGF Affibody-13 exhibited a 10-fold higher equilibrium dissociation constant than PDGF Affibody-16 on BLI. Moreover, hydrogels containing the original PDGF affibody and PDGF Affibody-11 displayed faster PDGF release despite their interactions with PDGF on BLI yielding equilibrium dissociation constants that were similar to that of PDGF Affibody-16. Thus, the effect of PDGF-specific affibodies on PDGF release could not be explained by the equilibrium dissociation constants alone.

These unexpected outcomes could be due to differences in PDGF-affibody binding within the crowded environment of the hydrogel polymer network compared to BLI binding interactions measured in solution. Interestingly, the PDGF release profile from hydrogels containing the original PDGF affibody was more consistent with the expected release profile from the equilibrium dissociation constant measured using yeast surface display, suggesting that the binding interaction measurements performed on yeast surface display may be more representative of binding within the hydrogel environment. Overall, these experiments demonstrate that rationally designed PDGF-specific affibodies could successfully control PDGF release from hydrogels at different release rates.

### VEGF-specific Affibodies Inhibit VEGF-induced HUVEC Proliferation

Computational modeling of binding interactions between VEGF and PDGF and their respective affibodies predicted binding at epitopes on the growth factors that were involved in receptor binding (**Fig. S1A, Fig. S2A**), suggesting a possible role of VEGF-specific and PDGF-specific affibodies in modulating protein bioactivity. As such, we next sought to investigate the effect of affibody-protein binding on protein bioactivity and downstream cellular responses.

VEGF bioactivity was evaluated using a human umbilical vein endothelial cell (HUVEC) proliferation assay, leveraging the well-documented ability of VEGF to prolong endothelial cell survival through a protein kinase C-dependent Raf-MEK-MAP kinase pathway (**Fig. 4A**).^16,56,57^ HUVECs were treated with 0.20-800 ng/mL of VEGF in reduced media for 96 hours to construct a VEGF dose-response curve. Cell proliferation was assessed using the CellTiter-Glo assay, in which metabolically active cells are detected via a luminescent signal caused by ATP-dependent luciferase activity. Luminescence increased with increasing VEGF concentrations in a dose-dependent manner, exhibiting a sigmodal dose-response curve with a maximum response measured at approximately 36,000 relative luminescent units (RLU) and a 50% effective dose (ED_50_) of 7.64 ng/mL of VEGF (**Fig. 4B**).

**Figure 4.**
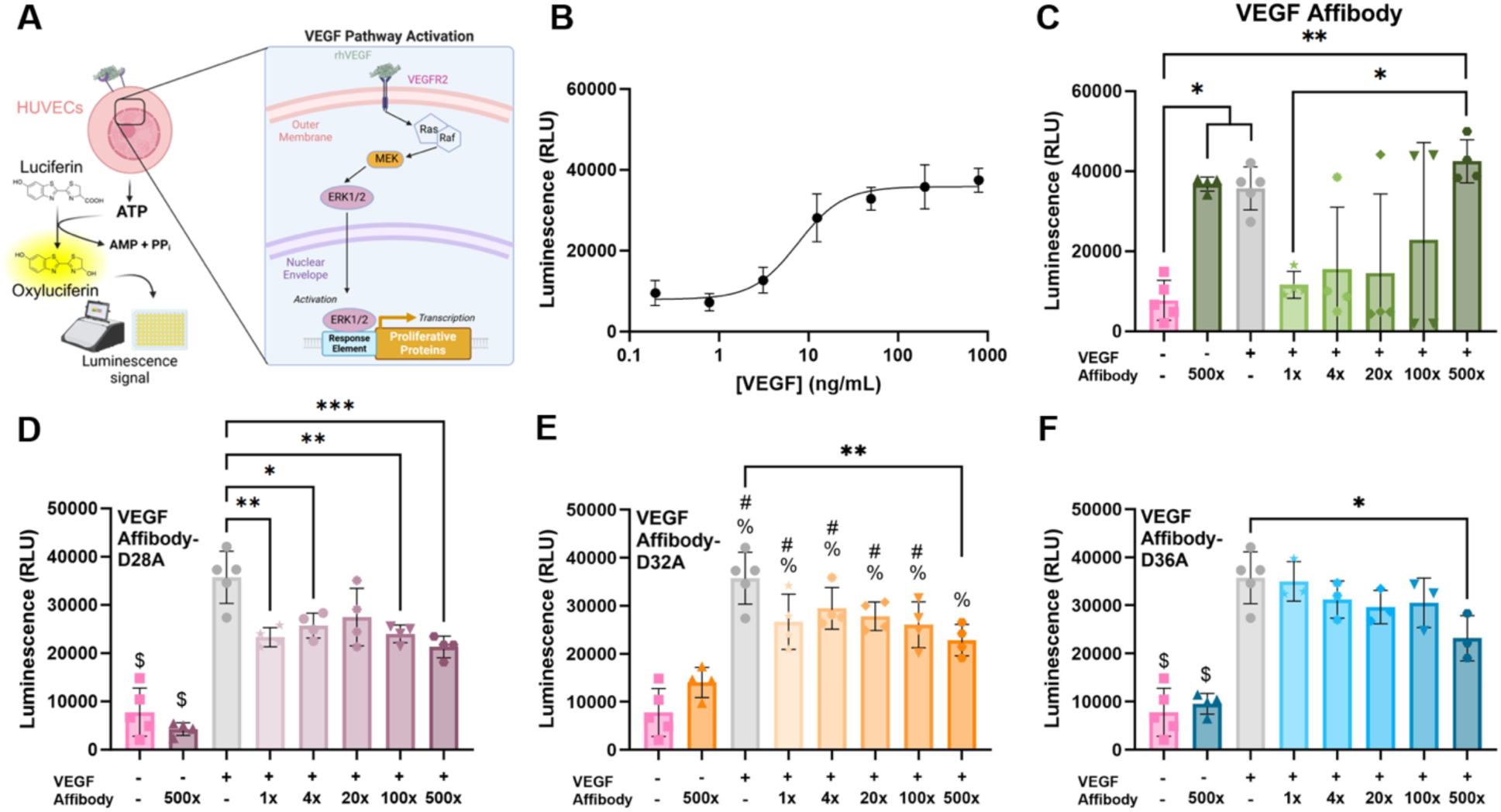
Effect of VEGF-specific affibodies on VEGF-induced proliferation of HUVECs. A) Schematic of VEGF-induced ERK1/2-dependent HUVEC proliferation and detection via a luminescent cell viability assay. B) Luminescence response curve of HUVECs treated with different concentrations of VEGF for 96 hours, demonstrating an EC_50_ = 7.64 ng/mL and R^2^ = 0.91. Response curve fit to a 4-parameter logistic model with 5 replicates at each concentration. HUVECs were treated with 19,100 ng/mL of affibodies (500 times molar excess to VEGF), 200 ng/mL of VEGF, or 200 ng/mL of VEGF pre-incubated with 1, 4, 20, 100, or 500 times molar excess of C) the original VEGF-specific affibody, D) VEGF Affibody-D28A, E) VEGF Affibody-D32A, or F) VEGF Affibody-D36A. Statistical significance was determined by one-way ANOVA and Tukey’s post-hoc test. (n = 4, *p < 0.05, **p < 0.01, and ***p < 0.001 as indicated. $ denotes a significant difference from all other groups. % denotes a significant difference from no treatment. # denotes a significant difference from 500 times molar excess of affibody.)

To determine the effect of VEGF-specific affibodies on VEGF bioactivity and subsequent VEGF-mediated HUVEC proliferation, 200 ng/mL of VEGF were pre-incubated with either 1, 4, 20, 100, or 500 times molar excess of VEGF-specific affibodies for 1 hour to allow VEGF-affibody binding to reach equilibrium. HUVECs were then incubated with the following treatments for 96 hours prior to luminescence detection: minimal media, 200 ng/mL of VEGF, 19,100 ng/mL of each VEGF-specific affibody (500x molar excess to VEGF), and the different molar ratios of each VEGF-specific affibody pre-incubated with VEGF. Unexpectedly, the original VEGF-specific affibody induced HUVEC proliferation in the absence of VEGF that was comparable to treatment with 200 ng/mL of VEGF (**Fig. 4C**). However, when VEGF was pre-incubated with an equal molar amount of the original VEGF-specific affibody, HUVEC proliferation was inhibited to comparable levels to no VEGF treatment. HUVEC proliferation increased up to comparable levels as treatment with 200 ng/mL of VEGF as the molar excess of the original VEGF affibody increased to 500x molar excess of VEGF affibody. In contrast, the mutant VEGF-specific affibodies did not induce HUVEC proliferation in the absence of VEGF and inhibited VEGF-induced HUVEC proliferation to different degrees when co-incubated with VEGF (**Fig. 4D-F**). Except for the pre-incubation condition of VEGF with 500x molar excess of VEGF Affibody-D32A, all groups treated with VEGF in the presence or absence of the VEGF-specific affibody mutants demonstrated significant increases in HUVEC proliferation compared to no treatment. For VEGF Affibody-D28A, pre-incubation of VEGF with affibodies inhibited HUVEC proliferation at all doses except for 20x molar excess of affibody. HUVECs treated with VEGF pre-incubated with VEGF Affibody-D32A and VEGF Affibody-D36A only inhibited HUVEC proliferation at the highest affibody dose of 500x molar excess to VEGF. Taken together with the BLI data, affibodies with higher affinities for VEGF may have a greater ability to inhibit VEGF activity, as the affibody mutant with the highest affinity for VEGF (VEGF Affibody-D28A) inhibited VEGF activity at the lowest doses. Similar results have been observed by others, in which rationally designed receptor inhibitors with higher affinities for their receptors more strongly inhibited downstream cellular responses.^39^

### PDGF-Specific Affibodies Modulate Downstream PDGFRβ Signaling

PDGF bioactivity was assessed using NIH/3T3 cells transfected with a firefly luciferase gene reporter plasmid that links multiple ERK1/2 signaling pathways, including PDGFRβ signaling, to luciferase expression through a serum response element under the regulation of ERK1/2 (**Fig. 5A**).^58,59^ Stably transfected NIH/3T3-Luc cells were seeded in serum-rich growth conditions for 12-16 hours, serum-starved for 24 hours, and then incubated with 0.195-12.5 ng/mL of PDGF for 5 hours. Firefly luciferase expression was assessed via the conversion of 5’-fluoroluciferin to oxyfluoroluciferin, which generates a stable luminescent output correlated to firefly luciferase concentration.^58^ Luminescence increased with increasing PDGF concentration in a dose-dependent manner, exhibiting a sigmoidal dose response with a EC_50_ of 2.94 ng/mL and maximal response of 23,655 RLU (**Fig. 5B**).

**Figure 5.**
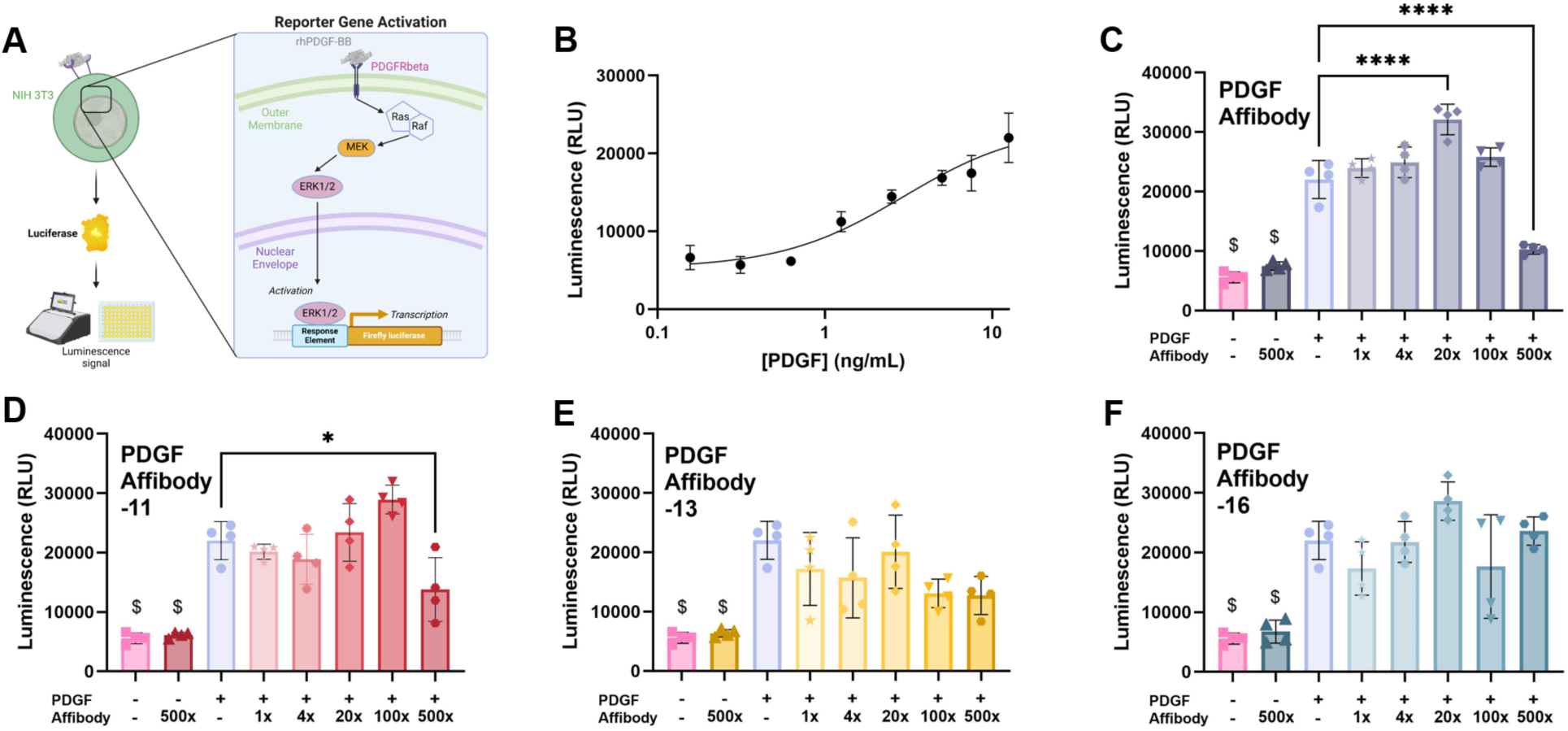
Effect of PDGF-specific affibodies on PDGF bioactivity measured using a PDGF-responsive cell line. A) Schematic of the NIH/3T3-Luc PDGF signaling pathway. NIH/3T3 cells were transfected with the luciferase reporter plasmid pGL4.33[luc2P/SRE/Hygro], which contains a serum response element that regulates luciferase expression as a function of ERK1/2-regulated transcriptional activity. B) Luminescence response curve of NIH/3T3-Luc cells treated with different concentrations of PDGF for 5 hours, demonstrating an EC_50_ = 2.94 ng/mL and R^2^ = 0.924. Response curve fit to a 4-parameter logistic model with 4 replicates at each concentration. NIH/3T3-Luc cells were treated with 1890 ng/mL of PDGF-specific affibodies (500 times molar excess to PDGF), 12.5 ng/mL of PDGF, or 12.5 ng/mL of PDGF pre-incubated with 1, 4, 20, 100, and 500 molar excess of C) the original PDGF-specific Affibody, D) PDGF Affibody-11, E) PDGF Affibody-13, or F) PDGF Affibody-16. Statistical significance was determined by one-way ANOVA and Tukey’s post-hoc test. (n = 4, *p < 0.05, ****p < 0.0001. $ denotes a significant difference from all other groups.)

To investigate the effect of PDGF-specific affibodies on PDGF bioactivity and PDGFRβ signaling, 12.5 ng/mL of PDGF was pre-incubated with 1, 4, 20, 100, or 500 times molar excess of PDGF-specific affibodies for 1 hour to allow PDGF-affibody binding to reach equilibrium. Pre-incubated PDGF and affibodies were added to NIH/3T3-Luc cells for 5 hours to induce PDGFRβ signaling and subsequent firefly luciferase expression. Treatment with any of the PDGF-specific affibodies alone did not induce luciferase expression, indicating that the PDGFRβ signaling cascade was not activated by the affibodies (**Fig. 5C-F**). NIH/3T3-Luc cells treated with PDGF pre-incubated with 500x molar excess of the original PDGF-specific affibody and PDGF Affibody-11 displayed significantly lower luminescence than treatment with the same concentration of PDGF alone, suggesting that these PDGF-specific affibodies inhibited PDGFRβ signaling. In contrast, no inhibition of PDGFRβ signaling was observed for PDGF pre-incubated with PDGF Affibody-13 (**Fig. 5E**) or PDGF Affibody-16 (**Fig. 5F**).

These results are generally consistent with the BLI data, in which PDGF Affibody-13 has the lowest affinity for PDGF and thus may have a limited ability to inhibit PDGFRβ signaling. Interestingly, a significant increase in luminescent signal was observed when PDGF was pre-incubated with 20x molar excess of the original PDGF-specific affibody compared to treatment with PDGF alone. This may have been due to the PDGF-affibody binding interaction stabilizing PDGF in a more favorable conformation for PDGFRβ binding. A similar effect has been observed with 14-3-3 sigma, known for its role in binding to phosphorylated insulin receptor 2 and generating a stable bound structure that causes prolonged downstream signaling through phosphoinositide 3-kinase.^60^

Collectively, the results of the HUVEC and NIH/3T3 experiments demonstrate that VEGF-specific and PDGF-specific affibodies have a range of inhibitory and stimulatory effects on VEGF and PDGF signaling, respectively, supporting the computational modeling results that suggest that these affibodies may bind to the VEGFR-2 and PDGFRβ binding epitopes on the proteins to modulate cell signaling.

### VEGF and PDGF Released from Affibody-Conjugated Hydrogels Retain Bioactivity

Both natural and engineered affinity-based interactions can prolong the stability and bioactivity of proteins released from biomaterials.^23,61,62^ Thus, we next explored the potential of our affibodies to prolong the bioactivity of proteins released from affibody-conjugated hydrogels. Either VEGF or PDGF was released from affibody-conjugated hydrogels with and without affibodies over 7 days into minimal media, protein concentrations were measured via ELISA, and released VEGF and PDGF were incubated with HUVECs or NIH/3T3-Luc cells to evaluate protein bioactivity at each timepoint. Cell responses were normalized to amount of VEGF or PDGF released at each time point to determine the specific activity of the released protein.

As expected from their relative affinities, hydrogels containing the original VEGF affibody released the least VEGF at Days 1, 2, and 4 (**Fig. 6A**). All affibody-containing hydrogels released bioactive VEGF out to Day 7, with hydrogels containing the original VEGF affibody and VEGF Affibody-D28A exhibiting higher specific activity at Day 4 (**Fig. 6B**). While hydrogels without affibodies released a nominal amount of bioactive VEGF over 7 days, the cumulative activity of VEGF released from all affibody-conjugated hydrogels was higher than VEGF released from PEG-mal hydrogels (**Fig. 6C**), suggesting that affibodies may stabilize and prolong VEGF bioactivity.

**Figure 6.**
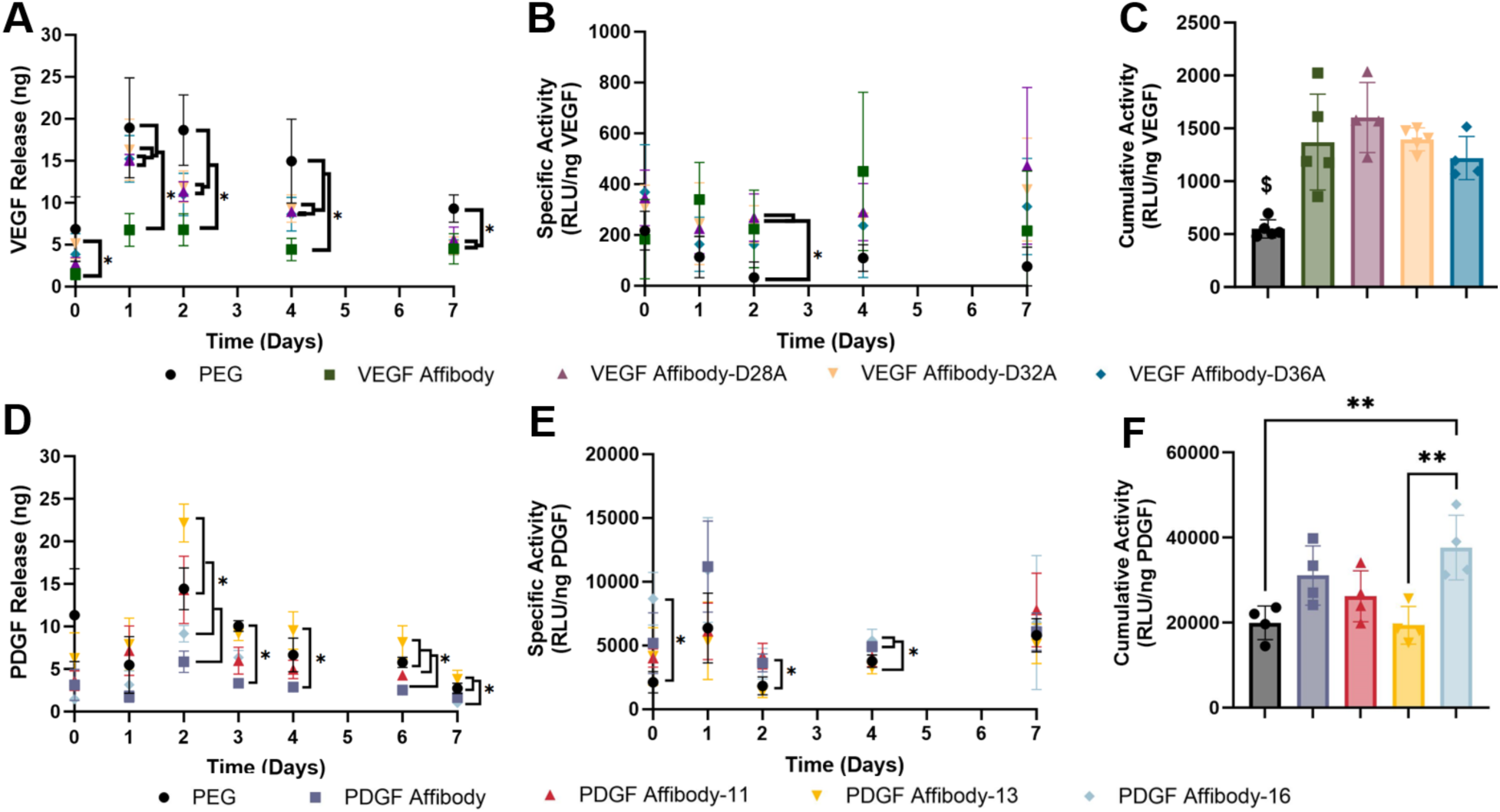
Bioactivity of VEGF and PDGF released from affibody-conjugated hydrogels. A) VEGF release into minimal media over 7 days from PEG-mal hydrogels without affibodies or containing the original VEGF affibody, VEGF Affibody-D28A, VEGF Affibody-D32A, or VEGF Affibody-D36A. Statistical significance was determined by two-way ANOVA and Tukey’s post-hoc test. (n = 4, *p < 0.05) B) VEGF-induced HUVEC proliferation normalized to amount of VEGF released at each timepoint. Statistical significance was determined by two-way ANOVA and Tukey’s post-hoc test. (n = 4, *p < 0.05) C) Cumulative bioactivity of VEGF released from hydrogels. Statistical significance was determined by one-way ANOVA and Tukey’s post-hoc test. ($ denotes a significant difference from all other groups.) D) PDGF release into minimal media over 7 days from PEG-mal hydrogels without affibodies or containing the PDGF original affibody, PDFG Affibody-11, PDGF Affibody-13, or PDGF-Affibody-16. Statistical significance was determined by two-way ANOVA and Tukey’s post-hoc test. (n = 4, *p < 0.05) E) Luminescence of PDGF-responsive NIH/3T3-Luc cells normalized to amount of PDGF released at each timepoint. Statistical significance was determined by two-way ANOVA and Tukey’s post-hoc test. (n = 4, *p < 0.05) F) Cumulative bioactivity of PDGF released from hydrogels. Statistical significance was determined by one-way ANOVA and Tukey’s post-hoc test. (n = 4, *p < 0.05, **p < 0.01)

Between Days 2 and 7, hydrogels without affibodies and hydrogels containing PDGF Affibody-13 or PDGF Affibody-16 intermittently released significantly more PDGF than hydrogels containing the original PDGF affibody (**Fig. 6D**). All hydrogels released bioactive PDGF for 7 days (**Fig. 6E**). Although hydrogels containing PDGF Affibody-16 released the least PDGF initially, this PDGF exhibited higher specific activity than PDGF released from hydrogels without affibodies. Despite a drop in PDGF activity at Day 2 across all groups, a steady increase in the specific activity of PDGF was observed between Days 2 and 7 for all hydrogels, suggesting the release of additional bioactive PDGF. Interestingly, PDGF released from hydrogels containing the original PDGF affibody, PDGF Affibody-11, and PDGF Affibody-16 displayed higher bioactivity at Days 2 and 4 compared to hydrogels without affibodies and hydrogels containing PDGF Affibody-13. Moreover, hydrogels containing PDGF Affibody-16 displayed higher cumulative PDGF activity over 7 days than hydrogels without affibodies and hydrogels containing PDGF Affibody-13 (**Fig. 6F**). The low cumulative bioactivity of PDGF released from PDGF Affibody-13 conjugated hydrogels may have been due to its significantly higher equilibrium dissociation constant, suggesting a relationship between the strength of the affinity interaction and prolonged protein bioactivity. Collectively, these results may suggest a stabilizing effect of PDGF-specific affibodies on PDGF bioactivity in hydrogels.

## Conclusion

Engineering specific protein-protein affinity-based interactions for controlling protein release can expand our ability to mimic the sustained presentation of proteins observed during tissue development and repair processes such as angiogenesis. However, engineering highly specific protein binders with tunable affinities and specificity between structurally similar proteins remains challenging and time-intensive using current approaches. Here, we report a library of new VEGF-specific and PDGF-specific protein binders as well as a computational pipeline for efficiently engineering highly specific protein binders from known starting scaffolds for use in affinity-controlled protein delivery vehicles.

Computationally informed point mutations disrupted VEGF-affibody interactions, generating mutants with weaker affinities for VEGF, association rate constants spanning five orders of magnitude, and significant differences in cumulative VEGF release from affibody-conjugated hydrogels. Additional PDGF-specific affibodies designed using Rosetta-based rational design demonstrated improved specificity towards PDGF compared to the initial PDGF-specific affibody. Unexpectedly, the PDGF-specific affibodies designed to exhibit higher affinities for PDGF demonstrated a range of binding kinetics that produced different PDGF release profiles from affibody-conjugated hydrogels.

Both soluble VEGF-specific and PDGF-specific affibodies affected downstream cellular responses of their target proteins, providing indirect evidence to support the computationally predicted binding interfaces at receptor binding epitopes and another method to modulate protein presentation and bioactivity. Finally, we demonstrated that affibody-conjugated hydrogels could sustain the bioactivity of VEGF and PDGF released into solution over 7 days at physiological temperature, providing a means to prolong protein bioactivity in therapeutic applications.

Overall, our work provides a promising platform for tuning VEGF and PDGF bioactivity and delivery, enabling future investigation into the temporal effects of these proteins in angiogenesis. The success of our computational approach for tuning the affinities of known VEGF-specific and PDGF-specific affibodies while retaining and improving target specificity indicates a promising avenue for accelerating the discovery of affinity-based protein-protein interactions with user-defined characteristics for controlled protein delivery.

## Acknowledgements

We are grateful to members of the Hettiaratchi lab for their thoughtful review of this manuscript. This work was supported by an Oregon Health & Science University (OHSU) Medical Research Foundation New Investigator Grant, a R21 Trailblazer Award and R35 Maximizing Investigators’ Research Award from the National Institutes of Health (NIH) (R21EB032112, R35GM147507), and a National Science Foundation (NSF) CAREER Award (2237240). J.E.S. is currently supported by an NIH Ruth L. Kirschstein Predoctoral Individual National Research Service Award (F31HL176164) and was previously supported by an NSF Research Traineeship in Molecular Probes and Sensors for Complex Environments (2022168). J.R.S. was previously supported by the University of Oregon Molecular Biology and Biophysics Training Program funded by the NIH (T32GM007759). M.R.F. was partly supported by the University of Oregon Summer Program for Undergraduate Research (SPUR), Knight Campus Undergraduate Scholars (KCUS) Program, and University of Oregon Vice President for Research and Innovation (VPRI) Undergraduate Fellowship. Figures were created using BioRender.com.

## Materials and Methods

### Materials

All reagents were purchased from Thermo Fisher Scientific or Sigma Aldrich, unless otherwise noted. Recombinant human VEGF165 and recombinant human PDGF-BB were purchased from PeproTech, biotinylated VEGF165 was purchased from Acro Biosystems, and biotinylated PDGF-BB was purchased from R&D Biosystems.

### Identification of VEGF- and PDGF-specific affibodies

To initially identify affibodies specific to VEGF or PDGF, cell sorting was performed on a yeast surface display library of the EBY100 strain of *Saccharomyces cerevisiae* containing approximately 4 × 10^8^ unique affibody sequences.^30^ Growth of the yeast surface display library and subsequent magnetic- and fluorescence-activated cell sorting steps were performed as previously described separately for each target protein.^34^ One VEGF-specific and one PDGF-specific affibody were selected for further analyses.

To prepare for computational modeling of VEGF binding interactions, a VEGF structure was derived from the high-resolution x-ray crystallography structure of the VEGF receptor binding domains (PDB: 2VPF)^63^ and pruned to remove all x-ray coordinates for co-factors and water atoms. The VEGF-VEGFR-2 binding interaction was modeled using elements from the known structure of VEGF interacting with VEGFR-2 (PDB: 3V2A) and the dimeric VEGF structure (2VPF).^64^ Alphafold2 (Google DeepMind) was used to generate a predicted structure from the sequence of the VEGF-specific affibody identified using yeast surface display.^53^ This structure had a predicted local distance difference test (pLDDT) score of greater than 95, indicating high confidence of the prediction. The computational modeling software ZDOCK was used to model the top 10 binding interfaces of the VEGF-affibody binding interactions.^43^ Docked structures were relaxed in Rosetta the to the lowest energy state, and scoring data were collected to select the bound structure with the greatest contact molecular surface area, shape complementarity, hydrophobic patch surface area, and polar contacts. In parallel, VEGF-affibody structures were compared to the known VEGF-VEGFR-2 binding interaction, and the binding interface with the greatest overlap with the known VEGFR-2 binding epitope on VEGF was selected as the most likely binding interface. Rosetta Docking confirmed the specificity of the original VEGF-specific affibody towards this single site on VEGF. This VEGF-affibody binding interface was visually inspected to observe the residues in direct contact between VEGF and the VEGF-specific affibody at the binding interface using PyMOL (Schrödinger). Three aspartic acids on the VEGF affibody at positions 28, 32, and 36 were predicted to participate in key polar contacts with residues on VEGF at the VEGF-affibody interface and individually mutated to alanine to disrupt the affinity-based interaction. Structures for each point mutant with pLDDT > 0.93 were generated using Alphafold2 and aligned at the predicted original VEGF affibody binding interface on VEGF to generate VEGF-affibody bound structures. Bound structures were relaxed and scored using Rosetta FastRelax prior to docking. Rosetta Docking was used to screen 5000 iterations of each mutant affibody binding across the surface of VEGF. Docking interface stability scores as a function of RMSD from the initial binding interface were plotted to determine the impact of mutagenesis on the binding interactions of each mutant affibody to VEGF. These three single point mutants were chosen for subsequent bacterial protein expression and characterization.

The x-ray crystallography structure of PDGF-BB complexed with PDGFR-β (PDB: 3MJG) was used to derive a PDGF structure for subsequent computational steps.^10^ Similar to the VEGF structure, the PDGF-PDGFR-β structure was pruned to remove all x-ray coordinates for co-factors, water atoms, and residues of the PDGFR-β, leaving only the PDGF x-ray coordinate data. Alphafold2 was used to generate a structure from the sequence of the PDGF-specific affibody identified using yeast surface display, which was predicted with high confidence (pLDDT > 93).^53^ The starting PDGF-affibody bound structure underwent energy minimization using Rosetta FastRelax scripts to globally relax surface-exposed residues of bound structure to the lowest energy states prior to docking.^42^ ZDOCK and HDOCK were applied in parallel to predict the 10 lowest energy interfaces to be created when the PDGF-specific affibody bound to PDGF at different possible binding sites.^43,44^ Physical characteristics of the PDGF-affibody bound structures from ZDOCK and HDOCK were quantified using Rosetta scoring scripts using ref2015.wts score function metrics and cross-referenced to select the interface with the greatest contact molecular surface area, shape complementarity, hydrophobic patch surface area, and polar contacts.^39,45^ The PDGF-specific affibody was predicted to form a stable binding interface on the PDGFR-β binding epitope of PDGF. Rosetta FastRelax was again applied to the PDGF-affibody complex to pack interfacial sidechains to their lowest energy conformations prior to designing new affibodies. Next, Rosetta FastDesign was used to mutate 18 affibody residues that were within 5 Å of the binding interface and to allow repacking of all PDGF and affibody residues within 8 Å of the binding interface, generating an *in silico* mutant library of 500 unique affibodies.^65^ Rosetta Score metrics were extracted from FastDesign run files to select the top 90th percentile of designed affibodies with binding interfaces that displayed high molecular contact surface area (cms > 400), zero buried unsatisfied polar contacts (vbuns_all < 0), low likelihood of binder aggregation (b_sap < 35), favorable predicted binding energy of the complex (ddG < -30), high hydrophobic surface area coverage upon binding (t_sap_score – tb_sap_score > 12), high shape complementarity (sc > 0.6), and stable structures (score < -70). 5000 rounds of Rosetta Docking scripts were run for each of these 10 PDGF-specific affibody candidates, modeling the affibody binding interface across the entire surface of PDGF with a step size of 0.5 Å RMSD from the starting designed binding interface at the PDGFR-β binding epitope. Interface stability scores as a function of RMSD from the initial binding interface were plotted to determine the specificity of the designed affibodies for the initial modeled interface between PDGF and the affibodies. Affibodies that exhibited a single, low-energy binding interface underwent an *in silico* folding stability screen using Robetta and Alphafold2 to determine if mutagenesis during FastDesign disrupted the alpha-helical folding of the affibody. Three PDGF-specific affibodies passed all *in silico* screening and were chosen for subsequent bacterial protein expression and characterization.

### Cloning affibody sequences into *E. coli*

VEGF and PDGF-specific affibody sequences were modified to contain a hexahistidine tag for protein purification and C-terminal cysteine for bioconjugation and codon-optimized for expression in *E. coli* using the Integrated DNA Technologies (IDT, Newark, NJ) codon optimization webtool. Each affibody-coding DNA sequence was inserted into the pet28b+ vector containing kanamycin resistance and the isopropyl-β-d-1-thiogalactopyranoside (IPTG)–inducible T7 promoter through restriction enzyme digestion and incubation with T4 ligase at 37 °C for 4 h. The plasmids were heat shock transformed into BL21 (DE3) *E. coli* (New England Biolabs, Ipswich, MA). *E. coli* were spread onto Luria-Bertani (LB) agar plates containing 50 µg/mL kanamycin sulfate and incubated for 16 hours at 37 °C. Single colonies were swabbed to inoculate liquid cultures of LB (Affymetrix, Cleveland, OH) with 50 µg/mL kanamycin sulfate for plasmid extraction and subsequent whole plasmid sequencing (Plasmidsaurus, Eugene, OR), followed by expansion for storage in 25% (v/v) glycerol at -80 °C.

### Soluble protein expression in *E. coli*

Small-scale bacterial culture was performed from single *E. coli* colonies to verify protein expression as previously described.^66^ For large-scale protein expression, 20 mL of LB supplemented with 50 μg/mL of kanamycin sulfate were inoculated with transformed *E. coli*. Cultures were incubated at 37 °C with orbital shaking at 250 rpm for 12-16 hours. Concurrently, 85.7 g of Terrific Broth (TB) powder (Research Products International, Mount Prospect, IL) and 0.4% (w/v) glycerol were dissolved in 1.8 L of ddH_2_O and sterilized by autoclaving. The following day, the 20 mL *E. coli* cultures were transferred to the TB supplemented with 50 μg/mL of kanamycin sulfate and 12-15 drops of antifoaming agent (Antifoam 204; Sigma-Aldrich). TB cultures were placed in a LEX-10 water bath bioreactor (Epiphyte, Toronto, ON, Canada) at 37 °C and aerated with lab air via gas sparger until an optical density at 600 nm (OD_600_) ≥ 0.7 absorbance units was reached, at which point protein expression was induced at 18 °C for an additional 14-18 hours via addition of 0.5 μM of IPTG (GoldBio, St. Louis, MO). Bacterial cultures were then centrifuged at 6,000 rpm and 4 °C for 20 minutes, and cell pellets were frozen at -80 °C.

For protein purification, cell pellets (5-10 g) were thawed in approximately 35 mL of binding buffer (50 mM Tris (GoldBio), 500 mM NaCl, 5 mM imidazole and 8.6 mM tris(2-carboxyethyl)phosphine HCl (TCEP; GoldBio)), sonicated on ice at 55% amplitude for 5 minutes (15 seconds on, 50 seconds off), and centrifuged at 13,000 rpm and 4 °C for 30 minutes. The supernatant was collected and agitated at 4 °C with 1.8 mL cobalt agarose beads (GoldBio) for a minimum of 4 hours. The mixture was poured into a glass chromatography column (BioRad) and washed with 5 x 10 mL of wash buffer containing TCEP (50 mM Tris, 500 mM NaCl, 30 mM imidazole, 10 mM TCEP) followed by 5 x 10 mL of wash buffer without TCEP. The protein was then eluted using elution buffer (50 mM Tris, 500 mM NaCl, 250 mM imidazole) in 1 mL increments until protein was no longer detected in the flow-through using Bradford reagent. The eluted protein was buffer-exchanged into phosphate buffered saline (PBS, Research Products International) and concentrated to 1-5 mg/mL using a 3 kDa molecular weight cut off (MWCO) centrifugal filter (Millipore, Burlington, MA). For further purification, size exclusion chromatography (SEC) was performed using a HiPrep 16/60 200 HR chromatography column (Cytiva, Marlborough, MA) on an NGC Chromatography System (Bio-Rad Laboratories, Hercules, CA). Affibodies were loaded into the column equilibrated with PBS. 5-10 consecutive 1 mL fractions were collected based on 280 nm absorbance signal and time of elution. Samples from each fraction were resuspended in 4x Laemmli dye for sodium dodecyl sulfate– polyacrylamide gel electrophoresis (SDS-PAGE) followed by Coomassie Brilliant Blue staining (Bio-Rad Laboratories) to visualize protein bands and confirm affibody purity. Selected fractions were then pooled, concentrated using a 3 kDa MWCO centrifugal filter, and stored at -80 °C.

### Matrix-assisted laser desorption ionization–time of flight mass spectrometry (MALDI)

Matrix-assisted laser desorption/ionization time of flight (MALDI-TOF) mass spectrometry was performed on purified affibodies using a Bruker Smart LS system (Bruker, San Jose, CA) to determine product size distribution, as described previously.^67^ Affibodies were frozen at -20 °C overnight followed by lyophilization at -105 °C and 40 mTorr using a VirTis BenchTop Pro freeze dryer (SP Scientific, Stone Ridge, NY). Affibodies were reconstituted in 3% (v/v) acetonitrile in ddH_2_O with 0.1% (v/v) trifluoracetic acid (TFA) (Oakwood Chemical, Estill, SC) at 0.7 mg/mL. 10 mg/mL of MALDI matrix was prepared by dissolving α-cyano-4-hydroxycinnamic acid in 0.1% (v/v) TFA and 50% (v/v) acetonitrile in ddH_2_O. 1 µL of matrix and sample were deposited on a stainless steel MALDI target plate (Bruker). The spectrometer was calibrated using the Protein Calibration Standard I (4000-20000 Da range, Bruker) prepared similarly to the affibody samples. Sample spectra were averaged over 200 readings. Resulting spectra were normalized to the prominent signal peak.

### Circular dichroism

Circular dichroism was performed on purified affibodies using a Jasco J-815 spectropolarimeter (Jasco, Easton, MD) to discern secondary structure. Affibodies were buffer-exchanged using a 3 kDa MWCO centrifuge filter into 10 mM Tris pH 7.4, which was a pH value at least 0.5 units away from all predicted affibody isoelectric points, and diluted to 0.3 mg/mL. Samples were loaded into 0.1 cm quartz cuvettes (Starna Corp), following which high tension voltage and absorption spectra across 190-250 nm were taken in triplicate using a step size of 1 nm at room temperature. Absorption spectra were averaged and normalized to measurements of 10 mM Tris pH 7.4. Circular dichroism output units (mdeg) were converted into molar ellipticity and normalized to affibody molecular weight and concentration.

### Biolayer interferometry

Binding interactions between VEGF, PDGF, and soluble protein-specific affibodies were measured using a GatorPlus biolayer interferometer (BLI; GatorBio, Palo Alto, CA). For measuring binding between VEGF and VEGF-specific affibodies, streptavidin-functionalized probes (GatorBio) were pre-soaked in PBS containing 0.05% (w/v) Tween 20 (PBST) for 20 minutes before a baseline reading was taken for 180 seconds. Probes were loaded with 25 nM of biotinylated VEGF (bVEGF) in PBST for 300 seconds until an approximate wavelength shift of 0.5 nm was achieved. Loaded probes were submerged in PBST until the baseline wavelength reading stabilized (approximately 300 seconds). 3.125-1000 nM of soluble VEGF-specific affibodies serially diluted in PBST were associated to the bVEGF-loaded probes for 600 seconds. Probes were then submerged into PBST or 600 seconds to measure dissociation of the affibodies. For measuring binding between PDGF and PDGF-specific affibodies, nitrilotriacetic acid (Ni-NTA) functionalized glass probes (GatorBio) were similarly pre-soaked in PBST, then loaded with 200 nM of PDGF-specific affibodies in PBST for 300 seconds until an approximate wavelength shift of 0.5 nm was achieved. Loaded probes were submerged in PBST until the baseline wavelength reading stabilized (approximately 300 seconds). 1.563-50 nM of soluble PDGF serially diluted in PBST were associated to the affibody-loaded probes for 600 seconds. Probes were then submerged into PBST for 600 seconds to measure dissociation of PDGF. Measurements were also taken of bVEGF loaded onto the streptavidin-functionalized probes without affibodies, affibodies loaded onto Ni-NTA probes without PDGF, and 0.0625-50 nM of PDGF without affibodies loaded onto Ni-NTA probes to subtract background signal from the data. Binding curves were normalized to data from probes loaded with only affibodies and only growth factors using GatorOne software 2.10 (GatorBio).

BLI was also performed using streptavidin-functionalized glass probes to measure binding of affibodies to their off-target proteins. To evaluate binding between PDGF-specific affibodies and VEGF, probes were loaded with 25 nM of bVEGF in PBST and 1000 nM of PDGF-specific affibodies were allowed to associate. To evaluate binding between VEGF specific affibodies and PDGF, probes were loaded with 25 nM of bPDGF and 1000 nM of VEGF-specific affibodies were allowed to associate. 1000 nM of VEGF- or PDGF-specific affibodies were also associated to empty probes to subtract any non-specific binding to the streptavidin-functionalized probes.

Binding curves were fit to a global best-fit, non-linear regression model using GraphPad Prism 10.1.1 in which R^2^ > 0.97 to determine the equilibrium dissociation constant (K_D_), on-rate constant (k_off_), and off-rate constant (k_on_) of each binding interaction.

### Hydrogel fabrication

100 μL 5% (w/v) 4-arm polyethylene glycol maleimide (PEG-mal, 20 kDa, Laysan Bio, Arab, AL) hydrogels were synthesized in 2.0 mL low retention microcentrifuge tubes as previously described.^34,68^ PEG-mal was reconstituted in PBS at pH 7.4, and VEGF- and PDGF-specific affibodies (500:1 molar ratio of affibodies to growth factor) were conjugated to PEG-mal through a Michael-type addition of the affibody C-terminal cysteine to the maleimide. Affibody-conjugated PEG-mal was then crosslinked with 10 mg/mL dithiothreitol (DTT) and swelled overnight in PBS pH 7.4 at 4 °C with gentle rotation. The hydrogels were washed to remove excess DTT prior to loading with 20 µL of 5 ng/µL PDGF or VEGF in PBSA overnight at 4 °C. Following protein loading, the supernatant was recovered to determine protein encapsulation into the hydrogel. 900 µL of 0.1% (w/v) BSA in PBS were added to each microcentrifuge tube to begin protein release, and the hydrogels were incubated at 37 °C for 7 days. 200 µL samples were removed and replaced with 200 µL of fresh 0.1% (w/v) BSA in PBS immediately after starting the incubation (0 h) at the following timepoints: 15 minutes, 30 minutes, 1 hour, 3 hours, 6 hours, and 1-7 days. Enzyme-linked immunosorbent assays (ELISA, Peprotech, Rocky Hill, NJ) were used to quantify the amount of VEGF or PDGF released into the supernatant in each timepoint. Cumulative protein release normalized to the amount of encapsulated growth factor was plotted over time.

### VEGF bioactivity assay

Human umbilical vascular endothelial cells (HUVECs, ATCC, Manassas, VA) were seeded at 2,500 cells/cm^2^ and expanded in complete Endothelial Growth Medium (EGM; Lonza, Walkersville, MD) at 37°C and 5% CO_2_. At 70-80% confluency, HUVECs were trypsinized and plated in a 96-well plate at 9,375 cells/cm^2^. Cells were allowed to adhere for 6-12 hours before being rinsed with PBS. Treatments were administered in reduced media consisting of 30 parts Endothelial Basal Media (EBM; Lonza) and 1 part EGM. To establish the dose response curve for VEGF treatment, cells were treated with a concentration series of 0.20-800 ng/mL VEGF for 96 hours. Additional control groups included cells treated with reduced media only and reduced media without cells. After incubation, 50 µL of media was removed from each well and replaced with 50 µL of CellTiter-Glo detection buffer (Promega, Madison, WI). The plate was placed on an orbital shaker for 2 minutes in the dark, followed by an additional 10 min of incubation. Luminescence was measured using a Synergy Neo2 plate reader (Agilent Technologies, Santa Clara, CA). The luminescence of the reduced media control well was subtracted from each treatment well prior to data analysis.

To determine whether VEGF released from hydrogels retained its bioactivity, hydrogels were synthesized with or without VEGF-specific affibodies, then loaded with 100 ng of VEGF. VEGF was released into DMEM over 7 days, and 220 µL aliquots were removed and replaced with fresh DMEM at the same timepoints used to evaluate VEGF release. VEGF released immediately (0 hours) and at days 1, 2, 4, and 7 was quantified by ELISA. HUVECs were seeded and treated with 50 µL of released VEGF from each hydrogel condition for 96 hours. Luminescent signal from released VEGF at each timepoint was normalized to VEGF concentration measured by ELISA and graphed as specific activity. Total VEGF activity was calculated by summing the activity of VEGF across each timepoint.

### Generation of a PDGF-responsive NIH/3T3 cell line

NIH/3T3 fibroblast cells (ATCC) were thawed, seeded at 10,000 cell/cm^2^, and expanded in growth medium consisting of Dulbecco’s Modified Eagle’s Medium (DMEM, Cytiva) containing 10% (v/v) fetal bovine serum (R&D Systems, Minneapolis, MN). Cultures were routinely maintained at 37°C and 5% CO_2_ and passaged at 60-70% confluence. After three passages of expansion, NIH/3T3 cells were grown in Eagle’s Minimal Essential Medium (EMEM, ATCC) for a single passage prior to transfection. Transfection mixtures were prepared according to the manufacturer’s protocols. Briefly, 2500 ng of luciferase reporter plasmid pGL4.33[luc2P/SRE/Hygro] (Promega), which contains a serum response element that regulates luciferase expression as a function of Rhoa GTPase activation and multiple mitogen activated phosphorylated kinase (MAPK) pathways, was combined with 12.5 µL of Lipofectamine Plus™ reagent and 227.5 µL of Opti-MEM buffer (Thermo Fisher Scientific, Waltham, MA) and mixed for 10 minutes at room temperature.^59^ Concurrently, Lipofectamine Plus™ mix was prepared by combining 14.3 µL of Lipofectamine Plus™ solution with 271.7 µL of Opti-MEM buffer solution for 10 minutes. Lipofectamine Plus™ mix was then combined with Plus Reagent DNA mix at a 1:1 ratio and incubated for an additional 15 minutes. During this step, NIH/3T3 cells were trypsinized and seeded into 96-well plates at 40,000 cells/cm^2^ in 135 µl of EMEM. Cells were then transfected by adding 65 µL of Lipofectamine Plus Reagent DNA mix per well and incubated for 48 h at 37°C and 5% CO_2_. Individual wells were combined to establish a polyclonal population of transfected cells. Cells were expanded and passaged 4 times between 60-70% confluency in growth medium containing 200 µg/mL hygromycin (selective medium) to remove the transiently transfected population, thencontinually passaged in selective medium to maintain the stably transfected NIH/3T3 cell population. Stably transfected NIH/3T3 cells, hereafter referred to as NIH/3T3-Luc, were expanded and stored in liquid nitrogen at post-transfection passage 5 until use.

### PDGF bioactivity assay

NIH/3T3-Luc cells were seeded at 31,250 cells/cm^2^ and incubated in 100 µL of growth medium overnight in 96 well plates. Growth medium was aspirated, and cells were incubated in DMEM for serum starvation overnight. To establish the dose response curve for PDGF treatment, serum-starved cells were treated with a concentration series of 0.625-12.5 ng/mL of PDGF for 4-6 hours to stimulate luciferase expression. Following treatment, 50 µL of ONE-Glo Luciferase detection reagent (Promega) was added to each well and allowed to incubate 3 minutes in the dark. Luminescence was then measured using SpectraMax I3 plate reader (Molecular Devices, San Jose, CA).

To determine whether the PDGF released from hydrogels retained its bioactivity, hydrogels were synthesized with or without PDGF-specific affibodies, then loaded with 100 ng of PDGF. PDGF was released into DMEM over 7 days, and 220 µL aliquots were removed and replaced with fresh DMEM at the same timepoints used to evaluate PDGF release. PDGF released immediately (0 hours) and at days 1, 2, 3, 4, 6, and 7 was quantified by ELISA. NIH/3T3-Luc cells were seeded, serum-starved, and treated with 50 µL of released PDGF from each hydrogel condition for 4-6 hours. Luminescent signal from released PDGF from each timepoint was normalized to PDGF concentration measured by ELISA and graphed as specific activity. Total PDGF activity was calculated by summing the activity of PDGF across each timepoint.

### Statistical Analysis

BLI data are presented with 95% confidence intervals. All other data are presented as mean ± standard deviation. Unless otherwise described, all data were analyzed and graphed using GraphPad Prism version 10.1.1 (Boston, MA). Statistical significance was determined using one-way or two-way analysis of variance (ANOVA) followed by the appropriate post-hoc test. Assumptions of equal variances and Gaussian distributions were verified. P < 0.05 was considered statistically significant.

