## Supplemental Information for "Applying computational protein design to engineer affibodies for affinity-controlled delivery of vascular endothelial growth factor and platelet-derived growth factor"

**Supporting Information for Publication**

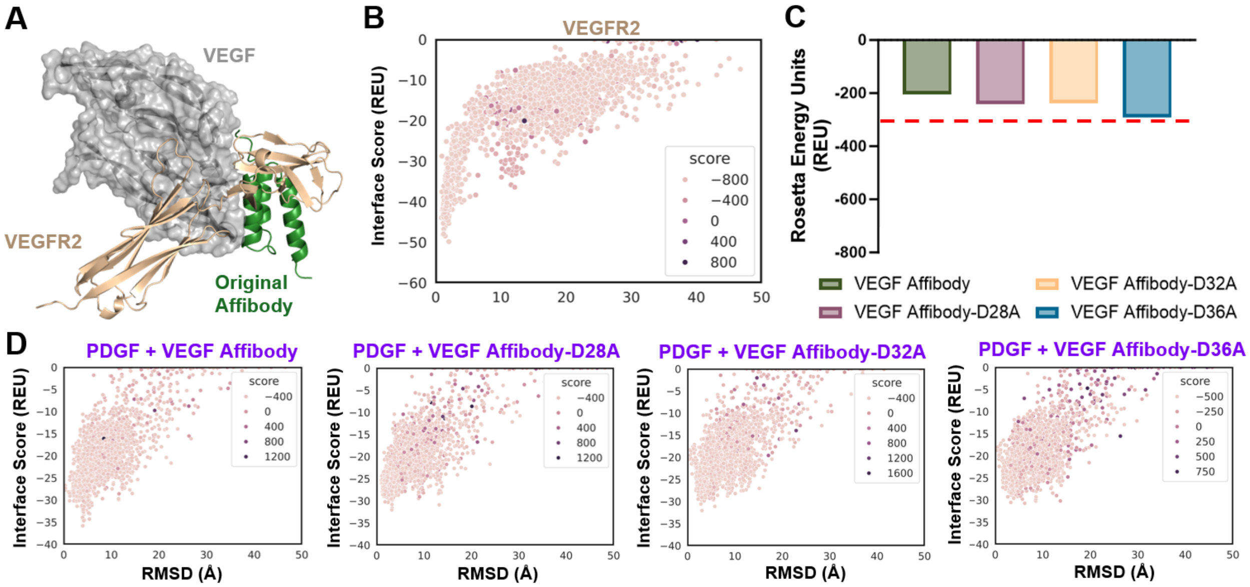


**Figure S1. Additional computational analysis of VEGF-VEGFR-2 binding and VEGF-specific affibodies binding PDGF.** A) Molecular modeling of the predicted original VEGF affibody binding site on VEGF compared to the known x-ray crystallography structure of VEGF bound to VEGF receptor-2 (VEGFR-2) (PDB ID: 3V2A). B) Rosetta Docking funnel depicting the interface stabilities of VEGFR-2 binding across the entire surface of VEGF (n=5000 docking interactions). C) Rosetta Score metrics measuring the predicted stability of VEGF-specific affibodies binding to PDGF. D) Rosetta Docking metrics measuring the predicted interface stability of VEGF-specific affibodies binding to PDGF.

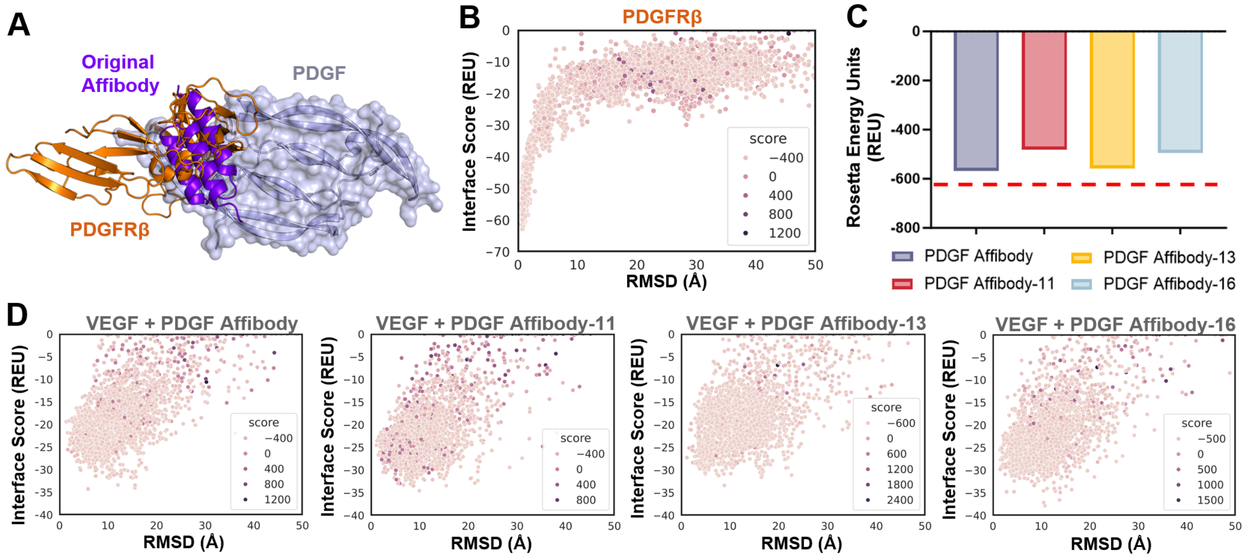


**Figure S2. Computational analysis of PDGF-PDGFRβ binding and PDGF-specific affibodies binding VEGF.** A) Molecular modeling of the predicted original PDGF affibody binding site on PDGF compared to the known x-ray crystallography structure of PDGF bound to PDGF receptor beta

(PDGFRβ) (PDB ID: 3V2A). B) Rosetta Docking funnel depicting the interface stabilities of PDGFRβ binding across the entire surface of PDGF (n=5000 docking interactions). C) Rosetta Score metrics measuring the predicted stability of PDGF-specific affibodies binding to VEGF. D) Rosetta Docking metrics measuring the predicted interface stability of PDGF-specific affibodies binding to VEGF.

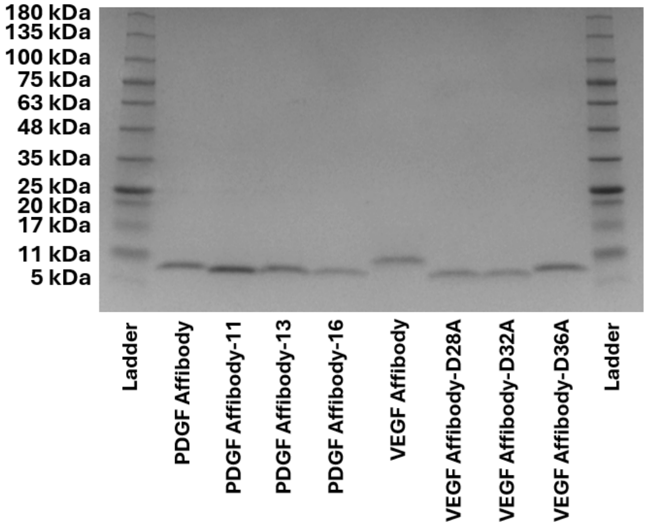


**Figure S3. SDS-PAGE gel of purified VEGF- and PDGF-specific affibodies.** Sodium dodecyl-sulfate polyacrylamide gel electrophoresis (SDS-PAGE) of affibodies and a 5-245 kDa reference ladder. Affibody samples were loaded at approximately 0.3 mg/mL. Gel was stained using Coomassie Brillant Blue.

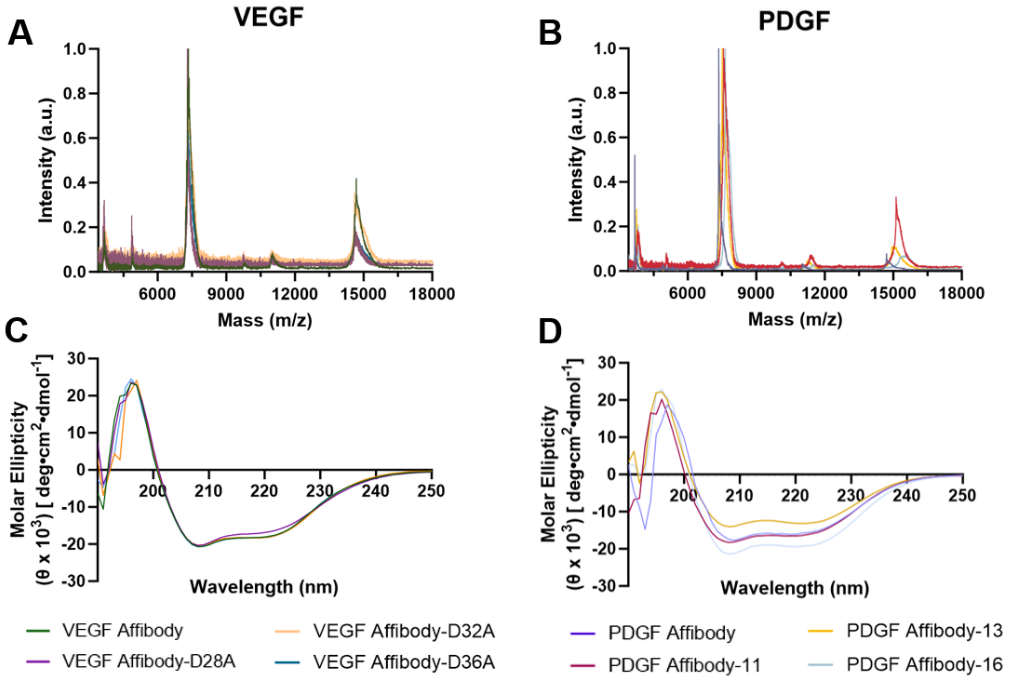

**Figure S4. Biochemical characterization of VEGF-specific and PDGF-specific affibodies.** Matrix-assisted laser desorption/ionization time of flight (MALDI-TOF) mass spectrometry spectra of A) VEGF-specific affibodies and B) PDGF-specific affibodies. Molar ellipticity of C) VEGF-specific affibodies and D) PDGF-specific affibodies measured over 190-250 nm using circular dichroism.

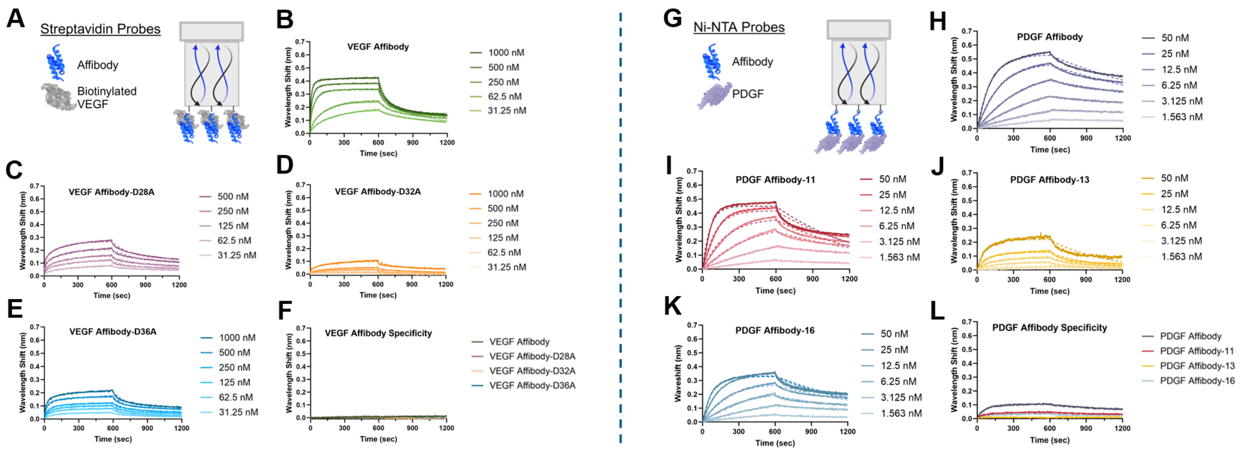


**Figure S5. Binding kinetics of VEGF-specific and PDGF-specific affibodies to VEGF and PDGF.** A) 25 nM of bVEGF were loaded onto streptavidin-coated BLI probes, followed by association and dissociation of 31.25-1000nM of B) the original VEGF-specific affibody, C) VEGF Affibody-D28A, D) VEGF Affibody-D32A, and E) VEGF Affibody-D36A. F) 25 nM of bPDGF were loaded onto streptavidin-coated BLI probes, followed by association and dissociation of 1000nM of each VEGF-specific affibody to evaluate binding specificity. G) 200 nM of PDGF-specific affibodies were loaded onto Ni-NTA BLI probes, followed by association and dissociation of 1.563-50 nM of PDGF. BLI results depicting binding interactions between PDGF and H) the original PDGF-specific affibody, I) PDGF Affibody-11, J) PDGF Affibody-13, and K) PDGF Affibody-16. L) 25 nM of bVEGF were loaded onto streptavidin-coated BLI probes, followed by association and dissociation of 1000nM of each PDGF-specific affibody to evaluate binding specificity. Solid lines depict measured data, while dashed lines depict fitted data used to generate binding kinetic constants.
